# Dynamic integration of cortical activity in the deep layer of the anterolateral superior colliculus

**DOI:** 10.1101/2024.07.20.604436

**Authors:** Hikaru Sugino, Sho Tanno, Tatsumi Yoshida, Yoshikazu Isomura, Riichiro Hira

**Author notes:** Corresponding authors. **Correspondence:** Riichiro Hira, M.D. Ph.D., Department of Physiology and Cell Biology, Graduate School of Medical and Dental Sciences, Tokyo Medical and Dental University, 1-5-45 Yushima, Bunkyo-ku, Tokyo, 113-8510, Japan. Yoshikazu Isomura, Ph.D., Department of Physiology and Cell Biology, Graduate School of Medical and Dental Sciences, Tokyo Medical and Dental University, 1-5-45 Yushima, Bunkyo-ku, Tokyo, 113-8510, Japan.

## Abstract

The superior colliculus (SC) receives inputs from various brain regions in a layer- and radial location-specific manner, but whether the SC exhibits location-specific dynamics remains unclear. To address this issue, we recorded the spiking activity of single SC neurons while photoactivating cortical areas in awake head-fixed Thy1-ChR2 rats. We classified 309 neurons that responded significantly into 8 clusters according to the response dynamics. Among them, neurons with monophasic excitatory responses (7–12 ms latency) that returned to baseline within 20 ms were commonly observed in the optic and intermediate gray layers of centromedial and centrolateral SC. In contrast, neurons with complex polyphasic responses were commonly observed in the deep layers of the anterolateral SC. Cross-correlation analysis suggested that the complex pattern could be only partly explained by an internal circuit of the deep gray layer. Our results indicate that medial to centrolateral SC neurons simply relay cortical activity, whereas neurons in the deep layers of the anterolateral SC dynamically integrate inputs from the cortex, SNr, CN, and local circuits. These findings suggest a spatial gradient in SC integration, with a division of labor between simple relay circuits and those integrating complex dynamics.

## Introduction

The superior colliculus (SC) has evolutionarily well-conserved vision-related functions. In mammals, it plays important roles in not only pure visual processing but also the intergration of movement-related visuospatial information (Wheatcroft, Saleem, and Solomon 2022; Isa et al. 2021; Gandhi and Katnani 2011; Wurtz and Albano 1980; Zahler et al. 2023, 2021; Takahashi, Sugiuchi, and Shinoda 2007; Ito, Feldheim, and Litke 2017; Sprague and Meikle 1965). The SC is also involved in other ethologically relevant behaviors, such as escape, defensive, foraging, and predatory behaviors (Gandhi and Katnani 2011; Dean, Redgrave, and Westby 1989; Basso and May 2017; Zingg et al. 2017; Shang et al. 2019; Wu et al. 2023; K. H. Lee et al. 2020; Li and Meister 2023). Furthermore, it plays an active role in higher-order brain functions that are thought to be primarily the responsibility of the cerebral cortex, such as attention, decision-making, and action selection (Müller, Philiastides, and Newsome 2005; Felsen and Mainen 2008; Comoli et al. 2012; Crapse, Lau, and Basso 2018; L. Wang et al. 2020; Duan, Pagan, et al. 2021; Essig, Hunt, and Felsen 2021; E. J. Jun et al. 2021; Thomas et al. 2023; Lovejoy and Krauzlis 2010; Stine et al. 2023; Glimcher and Sparks 1992). SC has also play roles in modulating learning in other brain regions such as the basal ganglia (Solié et al. 2022) and cerebellum (Kaku, Yoshida, and Iwamoto 2009; May, Warren, and Kojima 2024).

The SC has a distinct composition of seven alternating fibrous and cellular layers, which include three superficial (zonal [zo], superficial gray [SuG], and optic [op]), two intermediate (intermediate gray [InG] intermediate white [InWh]), and two deep layers (deep gray [DpG] and deep white [DpWh]) (Liu et al. 2022; May 2006). Although the superficial layer of the SC receives input from the retina and visual cortex, and thus its neural activity is highly selective for vision, the response properties of the intermediate to deep layer neurons activated by visual stimuli are linked to complex behaviors (Gharaei et al. 2020; Sparks and Hartwich-Young 1989; Ito, Feldheim, and Litke 2017; De Franceschi and Solomon 2020; González-Rueda et al. 2024). The intermediate and deep layers integrate multimodal information from the cerebral cortex, basal ganglia, and cerebellum (Miyashita and Tamai 1989; Komatsu and Suzuki 1985; Anderson and Yoshida 1977; Niemi-Junkola and Westby 2000). Specifically, inputs from the primary motor (M1) and secondary motor cortices (M2) are involved in the preparation and planning of movement and are important for the coordination of complex movements (Doykos et al. 2020; Duan, Pan, et al. 2021; Borra et al. 2014; Distler and Hoffmann 2015).

The SC can also be radially divided into distinct zones extending along the mediolateral (ML) axes. Although functional studies typically arbitrarily bisect the SC into a medial and lateral zone, a recent study identified four zone-specific medial-to-lateral delineations in the SC: medial SC, centromedial SC, centrolateral SC, and lateral SC, each of which possesses distinct circuitry within the cortico-tectal network (Benavidez et al. 2021). Henceforth, we will refer to the radially divided structure as ‘zone.’ The SC primarily receives direct projections from the visual cortex, whereas the centrolateral and lateral SC receive inputs from the frontal cortex. Additionally, the centrolateral SC receives projections from the posterior parietal cortex, whereas the lateral SC receives projections from higher-order cortical areas, such as the prefrontal cortex, which indicates the involvement of these zones in higher-order functions. The projections from the substantia nigra pars reticulata (SNr) and the output nucleus of the basal ganglia also have topography (Foster et al. 2021; Benavidez et al. 2021; McElvain et al. 2021).

Despite extensive studies on the anatomical segregation of layer and zone structures and the functional role in sensory integration and behavior output, it remains unclear what types of temporal dynamics occur at different SC locations. Therefore, by using ChR2-transgenic rats and a high-density silicon probe, Neuropixels (J. J. Jun et al. 2017), we recorded the evoked activity of single SC neurons in response to photoactivation of the dorsal cortex, including M1, M2, and S1 (Yoshida et al. 2024). We found that the temporal complexity of evoked activity depended on all layers, zones, and the anteroposterior (AP) location of the SC, and the stimulated area of the cerebral cortex. The anatomical basis for these different dynamics and how these particular dynamics contribute to various behaviors will also be discussed.

## Materials and methods

### Animal preparation

All experiments were approved by the Animal Research Ethics Committee of the Institutional Animal Care and Use Committee of Tokyo Medical and Dental University (A2019-274) and were carried out in accordance with the Fundamental Guidelines for Proper Conduct of Animal Experiment and Related Activities in Academic Research Institutions (Ministry of Education, Culture, Sports, Science and Technology of Japan). We used 6 adult male and female rats (Thy1.2-ChR2-Venus transgenic rats, Long–Evans strain (Tomita et al. 2009; Saiki et al. 2018) aged 10-14 weeks for SC recording. These rats were kept under an inverted light schedule (lights off at 12 AM; lights on at 12 PM) in their home cages to adapt to experimental surroundings.

### Surgery

Primary surgery was performed to attach a custom-made head-plate (Isomura et al. 2009) to the skull of rats under anesthesia with isoflurane gas (4.0–4.5% for induction and 2.0–2.5% for maintenance, Pfizer Japan, Tokyo, Japan) using an inhalation anesthesia apparatus (Univentor 400 anesthesia unit, Univentor, Zejtun, Malta). The body temperature was maintained at 37.0°C using an animal warmer (BWT-100, Bio Research Center, Aichi, Japan). The head of the rats was fixed on a stereotaxic frame (SR-10R-HT, Narishige) with ear bars, and lidocaine was applied (Xylocaine Jelly, Aspen Japan, Tokyo, Japan) for local skin anesthesia and povidone-iodine disinfectant solution (10%, Kaneichi, Osaka, Japan) for disinfection around surgical incisions. The head-plate (Isomura, Harukuni et al., 2009) was then glued to the skull with stainless steel screws and dental resin cement (Super-Bond C & B, Sun Medical, Shiga, Japan; Unifast II, GC Corp., Tokyo, Japan). Reference and ground electrodes (PFA-coated silver wires, A-M systems, WA; 125-mm diameter) were implantedunder the bone on the cerebellum. Analgesics and antibiotics (meloxicam, 1 mg/kg sc, Boehringer Ingelheim, Tokyo, Japan; gentamicin ointment, 0.1% us. ext., MSD, Tokyo, Japan) were finally applied to relieve pain and prevent infection. More than three days later, secondary surgery was performed under isoflurane anesthesia. For insertion of recording electrodes, we made a cranial window onto the cerebral cortex of the left hemisphere. The bone and dura mater were opened and removed by a dental drill (Tas-35LX, Shofu, Kyoto, Japan) and a dura picker (DP-T560-80, Bio Research Center, Aichi, Japan). The cortical surfaces were washed with PBS containing antibiotics (0.2% amikacin sulfate, Sawai, Osaka, Japan). We thinned the skull over the dorsal cerebral cortex and applied silicone oil to keep it transparent for optogenetic stimulation (**Fig. 1A**). The thinned-skull area covered M1, M2, S1, and PPC. Because the barrel cortex was not always completely covered with the thinned-skull window, we excluded the barrel cortex from S1 for the analysis when comparing the responsiveness. Analgesics (meloxicam, 1 mg/kg sc, Boehringer Ingelheim, Tokyo, Japan) was applied to relieve pain.

**Figure 1.**
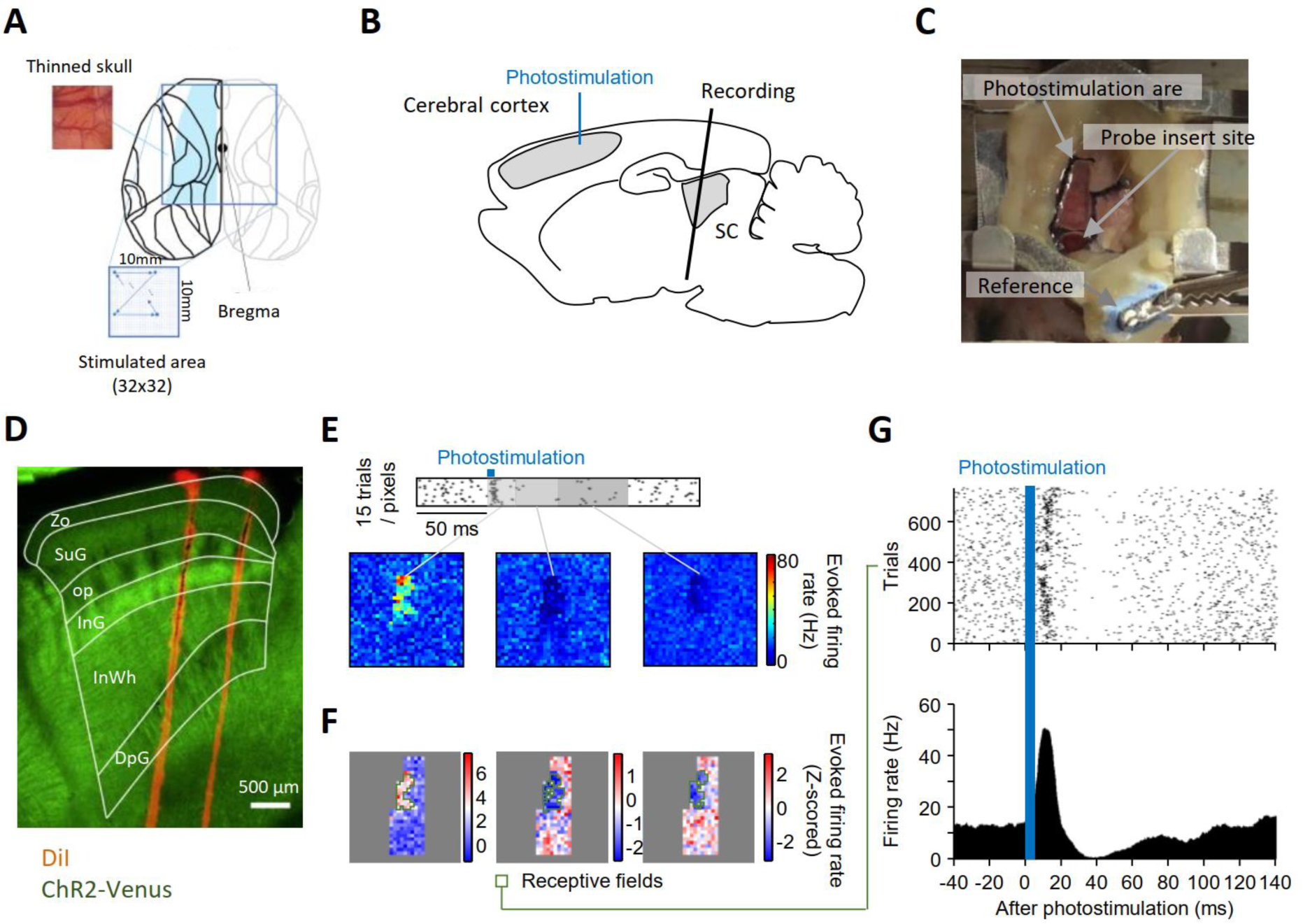
The experimental setup and basic analysis. (A) Dorsal view of the brain atlas. The blue square represents the 10 × 10 mm stimulation area. The black dot denotes the bregma. The blue-shaded area is the thin skull area. (B) Approximate positions of a recording site. (C) An aerial view before photostimulation of the cerebral cortex and recording of the SC. (D) A sagittal section of the SC in a ChR2-Venus transgenic rat. DiI fluorescence shows electrode location. White lines indicate the layer boundaries. (E) Colormaps of the mean firing rates evoked 0–20 ms (left), 20–50 ms (middle), and 50–100 ms (right) after photostimulation of each stimulation point. (F) Colormaps of Z-scored mean firing rates evoked 0–20 ms (left), 20–50 ms (middle), and 50–100 ms (right) after photostimulation of each stimulation point. The areas surrounded by a light green square correspond to the receptive fields. (G) The raster plot and PSTH relevant to the photostimulation of receptive fields. The blue bar corresponds to the photostimulation duration. SC, superior colliculus; Zo, zonal; SuG, superficial gray; op, optic; InG, intermediate gray; InW, intermediate white; DpG, deep gray

### Optogenetics

We used a custom-made scanning system for photoactivation of cortical neurons (Yoshida et al. 2024; Hira, Ohkubo, Tanaka, et al. 2013; Hira, Ohkubo, Ozawa, et al. 2013; Hira et al. 2015). For functional mapping, we stimulated a 32 x 32 grid (3.23 mm spacing, total 1,024 points) with 145 ms interval pseudo-randomly for 15 cycles. The stimulation point after the thinned-skull point was always at the bone, ensuring that the stimulation interval for the cerebral cortex was more than 290 ms. Recording during the stimulation at the bone was monitored as an internal control to discriminate with any unnecessary effects including the neuronal activity via retinal light responses. We confirmed that there was no contamination of visually evoked activity in the responsive neurons that we analyzed. Each stimulation was 5 ms duration with a 445 nm TTL-controllable diode laser (JUNO, Kyocera SOC, Japan). Laser power was set in each experiment sufficiently small not to induce movement when the primary motor cortex was stimulated (3.3 mW–3.9 mW at the center of field-of-view). Even at the edge of the field of view (7.1 mm from the center), the laser power was >85.7% of the center. All systems were controlled with custom-made software written with LabVIEW (National instruments, USA).

### Electrophysiology

Three hundred eighty-four channels in Neuropixels 1.0 (IMEC, Belgium) were used for simultaneous recording at a 20 μm step from the tip (total 7.6 mm long). Just before recording, the probes were manually coated with lipophilic dye, DiI (DiIC18(3), PromoKine, Heidelberg, Germany). The probes were fixed to a holder and handled using a micromanipulator (SM-15M, Narishige, Japan). Reference and ground electrodes were soldered, and attached to a custom-made part with four independent silver wires located under the skull onto the cerebellum. The probe was lowered in the coronal plane at 10 degrees from vertical at a speed of 10 μm/s until 8 mm below the dura for recordings of SC. They were left to settle for more than 30 min to reduce subsequent drift due to post-insertion brain relaxation. Recordings were controlled with Open Ephys software (version 0.5, https://github.com/open-ephys/plugin-GUI/releases) (Siegle et al. 2017). Spiking activity and local field potential (LFP) were obtained at 30,000 samples/s and 2,500 samples/s, respectively after 500 Hz high-pass filtering and band-pass filtering with 0.5–1kHz range, respectively. We recorded the timing and location of photostimulation at the same time with USB-6001 (National instruments, USA) to align timing of each photostimulation with the neural signals. We finally determined the location of each channel at the level of the subdivision within SC by histological observations. We recorded neurons from the SC.

### Analysis of electrophysiological data

All analyses were conducted with MATLAB (R2020a, R2022a, or R2024a, Mathworks). Spike sorting was performed using KiloSort3 (https://github.com/MouseLand/Kilosort). Sorted clusters were manually curated using Phy (https://github.com/cortex-lab/phy). The spontaneous firing rate of each neuron was obtained by averaging the firing rate during the 50 ms before each photostimulation. To identify “responsive neurons” to the stimulus of the cerebral cortex and “receptive fields” of each responsive neuron, we first extracted the candidate sections that satisfy the following criteria:

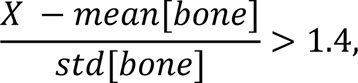

where the *X* value is the mean firing rate of the neuron 0–20, 20–50, or 50–100 ms after photostimulation of the section. The [bone] refers to stimulations performed at the skull surface.. We then excluded candidate sections surrounded by four or fewer candidate sections.. Finally, if there is any candidate section that is surrounded by five or more candidate sections that are also surrounded by five or more candidate sections among the eight surrounding sections, we defined the neuron and the candidate sections as responsive neuron and receptive fields, respectively. This criterion selects stimulated sections forming a local cluster, based on the premise that the distance between sections is smaller than the resolution of the stimulus. We confirmed that it identified fewer neurons than those were visually identifiable, allowing us to limit our analysis to neurons that had responded unambiguously. PSTHs were created by collecting the firing responses to the receptive fields from 50 ms before to 150 ms after the photostimulation in each trial. The peak latency was determined by finding the maximum or minimum value after convolving the PSTH with a Gaussian function with a standard deviation of 10 ms. To obtain average response maps, we aligned each map based on the bregma and the angle of midline before averaging. K-means clustering was preformed using a MATLAB function “kmeans”. We defined the excitatory response using the following criterion:

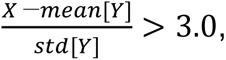

and inhibitory response using the following criterion:

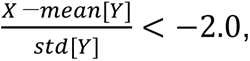

where *X* is the evoked firing rate at each time of 0–140 ms after photostimulation, and *Y* is the firing rate 0–40 ms before photostimulation. We then defined the duration of the excitatory response as the excitatory duration, the duration of the the inhibitory response as the inhibitory duration, and the sum of these durations as the total duration. Phases in neural responses were counted as the number of excitatory and inhibitory responses. However, if the response firing rate did not persist for at least 3 ms, it was not counted. Additionally, if the firing rate changed from an excitatory or inhibitory response to the basal firing rate but returned to the same firing rate within 3 ms, it was considered a continued response and counted as one phase.

K-means clustering was performed using a MATLAB function “kmeans”. Visual inspection of how the activity patterns were classified as the k value was increased revealed that mixed activity patterns in the same cluster were observed when k<8. Conversely, when k>8, clusters consisting of a small number of neurons frequently appeared. In the case of k=8, pairs that were initially in the same cluster remained in the same cluster 63.7% of the time (12.5% if random) when clustering was repeated 100 times with different initial conditions.

### SC classification into the subdivisions

We divided the SC into Zo, SuG, op, InG, InWh, DpG, and DpWh by the sagittal histological image of ChR2-Venus. We divided the SC into the medial zone, centromedial zone, centrolateral zone, and lateral zone radially by the coordinate (Benavidez et al. 2021). We divided the SC into three portions; the portion ≤ 5500 μm posterior to the bregma as anterior, the portion 5000–6000 μm posterior to the bregma as intermediate, and the portion ≥ 6000 μm posterior to the bregma as posterior.

### Histology

After all recording sessions, the rats were perfused transcardially with saline and subsequent 4% formaldehyde in 0.1 M phosphate buffer under deep anesthesia with urethane (3 g/kg, ip, Nacalai Tesque, Kyoto, Japan) The brains were removed and then post-fixed with the same fixative for at least 12 hours at 4° Celsius. The brains were removed and post-fixed with the same fixative for >2 days. The brains were then embedded in 2 % agar in PBS and cut into sagittal sections with a thickness of 50 µm using a microslicer (VT1000S, Leica, Wetzlar, Germany). The serial sections were cover-slipped with mounting medium (DAPI Fluoromount-G, Southern Biotech, AL, USA). DiI, Venus, DAPI, and Neuro Trace were visualized through a fluorescence microscope (IX83 inverted microscope, Olympus, Tokyo, Japan).

### Tracer experiment

Adult Thy1.2-ChR2-Venus transgenic rats (n = 7, male: 5, female 1, 180–400 g) were used in this experiment. The rats were anesthetized with 2.5 % isoflurane and an air flow of 0.2 liter per minute. Injection of AAVretro-CAG-tdTomato (addgene, Plasmid #59462-AAVrg, 2.4×10^12^ GC/ml, 0.2 µL, 0.02 µL/min or 0.5 µL, 0.05 µL/min, total 10 min) was performed (Hamilton, 10 µL Microliter Syringe Model #801 RN) at the VA complex (n=1, 1.6 mm left lateral to midline, 1.7 mm posterior to bregma, and 6.0 mm deep from the brain surface), VL (n=3, 1.7 mm left lateral to midline, 2.3 mm posterior to bregma, and 5.8∼6.1 mm deep from the brain surface), VM (n=2, 1.4 mm left lateral to midline, 2.4 mm posterior to bregma, and 6.9 mm deep from brain surface), or CL (n=1, 1.4 mm left lateral to midline, 2.8 mm posterior to bregma, and 5.2 mm deep from the brain surface) of thalamus by pressure through a glass micropipette. Two weeks after AAV injection, the rats were anesthetized with urethane, and then perfused intracardially with saline and fixative solution containing 4 % paraformaldehyde. The brain was quickly removed and fixed in 4% paraformaldehyde for 2 days at 4 °C then placed in a 30% sucrose in PBS solution for several days until they sank to the bottom of their container. The brain was then immediately immersed in isopentane over dry ice and then stored at −80 °C until sectioned. Sagittal sections (20 μm or 50 μm) were obtained in a cryostat (NX50, Epredia) at −20℃ through SC and thalamus. Some of the results in the other brain regions have already been shown in our previous work (Yoshida et al. 2024).

### Analysis of the tracer database

We used a public database of mouse connectome (Allen Mouse Brain Connectivity Atlas) (Oh et al. 2014). We selected two experiments where the anterograde trace, AAV2/1.pSynI.EGFP.WPRE.bGH, was injected in lateral SC (ID: 175158132) and centrolateral SC (ID: 128001349). The injection volume was 0.31 mm^3^ and 0.28 mm^3^, respectively. The injected location was AP: -3.8 mm, ML: 1.5 mm, DV: 2.1 mm, and AP: -4.36 mm, ML: 1.25 mm, DV: 1.2 mm relative to the bregma, respectively. We displayed the two-photon fluorescence imaging data of the injection site, thalamus, midbrain, brainstem, and cerebellar nuclei for each experiment.

### Statistics

Spearman’s rank correlation coefficient was used for pairwise comparison. Analysis of variance (ANOVA) was used for multiple-comparison. Bootstrap test, in which the values are calculated 10,000 times with shuffled data, and the proportion of shuffled values that are greater than or less than the actual value is taken as the p-value, was used for two or three dimensional data. Multiple regression analysis was performed using the four parameters; total duration, excitatory duration, inhibitory duration, and phase count, in each voxel as the dependent variable and the ML, AP, and DV coordinates of each voxel as the explanatory variables, and the lines passing through the center of mass of the plot and having each regression coefficient as the direction vector were displayed (**Fig.4C–F, K–N**). The F-test was used to determine the significance of the results of multiple regression, and a p-value of less than 0.05 was considered significant. Significance of each dependent variable was tested based on the *t*-value for each parameter. Data were expressed as means ± S.E.M unless otherwise noted.

## Results

### Classification of single SC neurons according to temporal activity patterns in response to cortical photoactivation

To stimulate large cortical areas in a point-by-point and compare the stimulated areas, we used transgenic rats (Saiki et al. 2018; Mitani et al. 2022) densely expressing ChR2 under Thy1 promoter in all layers of the cerebral cortex with their heads fixed in an awake state. We used a custom-made scanning system for the photoactivation of cortical neurons (Yoshida et al. 2024). We recorded the neural activity from the SC using a Neuropixels probe (J. J. Jun et al. 2017) (**Fig. 1A, B**, and **C**), isolating 758 neurons from six rats across 13 sessions. We identified the exact recording locations of each channel using histological data at the subdivision level (**Fig. 1D, Methods**). **Figures 1E and F** show the responses of an example neuron. The average firing rates at 0–20, 20–50, and 50–100 ms post-stimulus at each point are displayed on a color scale (response map). The response map indicated that the neuron received inputs from S1, M1, and M2. The stimulation points affecting the firing rate of the recording neuron were referred to as the ‘receptive field’ of the neuron. Raster plots and PSTHs for the receptive field of example neurons are shown in **Figure 1G**. This analysis was conducted for all neurons. We identified 309 ‘responsive neurons’ (40.1%) that had a receptive field (see **Methods**).

The SC neurons showed a variety of activity patterns in response to the stimulation of cortical areas. To characterize these temporal activity patterns, we conducted K-means clustering (K = 8) on the average responses to the photostimulation of the receptive field of all responsive neurons (**Fig. 2A; Methods**). This resulted in the classification of eight types of SC neurons: fast excitation (cluster #1: 13.3%, 41/309; cluster #2: 26.7%, 83/309; and cluster #3: 13.9%, 43/309), excitation with long duration (cluster #4: 7.4%, 23/309), triphasic excitation-inhibition-excitation (cluster #5: 10.7%, 33/309), inhibition followed by excitation with long duration (cluster #6: 13.3%, 41/309), inhibition (cluster 7: 7.8%, 24/309), and excitation followed by prolonged inhibition (cluster #8: 6.8%, 21/309; **Fig. 2B**).

**Figure 2.**
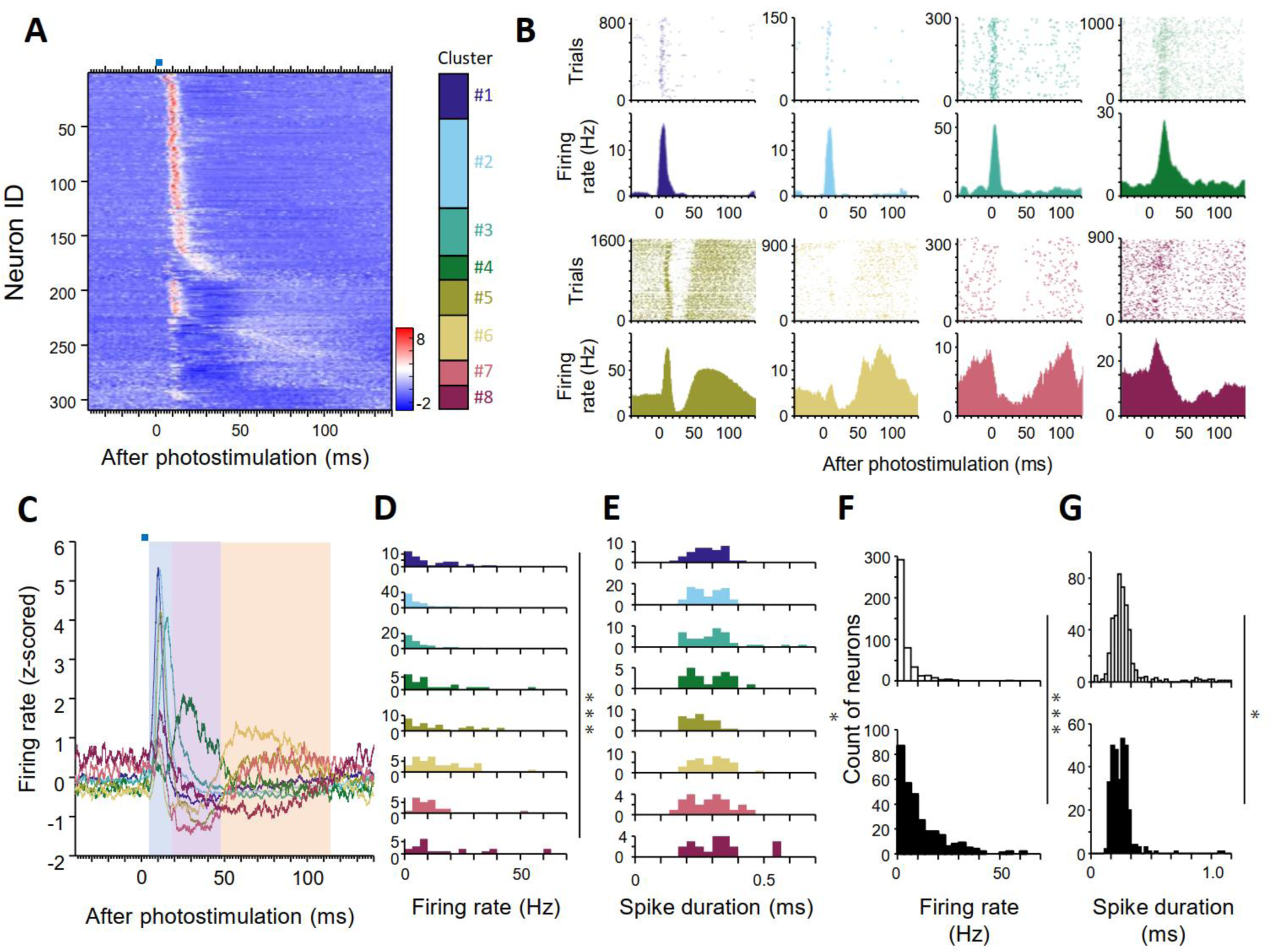
Classification of SC responses. (A) Z-scored mean firing rates of all responsive neurons. Color bar indicates the cluster number of each neuron: cluster #1 (indigo), cluster #2 (cyan), cluster #3 (teal), cluster #4 (green), cluster #5 (olive), cluster #6 (sand), cluster #7 (rose), and cluster #8 (wine). (B) Examples of raster plots and PSTHs of neurons classified into each cluster. Colors correspond to the cluster number in (A). (C) Mean z-scored activity of each cluster. Blue-, purple-, and orange-shaded areas correspond to periods 1, 2, and 3, respectively. (D) Baseline firing rates of neurons classified into each cluster. \*\*\**p* = 5.7×10^-8^, ANOVA. (E) Spike durations of neurons classified into each cluster. *p = 0.002, Wilcoxon’s rank-sum test. (F) Baseline firing rates of non-responsive neurons (white) and responsive neurons (black). \*\*\**p* = 4.5×10^-28^, Wilcoxon’s rank-sum test. (G) Spike durations of non-responsive neurons (white) and responsive neurons (black). \**p* = 0.0012, Wilcoxon’s rank-sum test.

To further characterize these clusters, we compared the firing rate and spike duration of the recorded neurons. Among the responsive neurons, the baseline firing rate varied between clusters (*p* = 7.2 × 10^-7^, ANOVA); specifically, clusters #4 and #5 had lower baseline firing rates than the other clusters (**Fig. 2D**). Notably, the distribution of spike widths appared to be bimodal in cluster #2, #3, #4, #6, #7, and #8 (**Fig. 2E**). The spike duration of cluster #5 was significantly shorter than that of other clusters (*p* = 0.002, Wilcoxon’s rank-sum test). However, multiple comparisons did not show significant differences between clusters (*p* > 0.05, ANOVA; **Fig. 2E**). The baseline firing rate was higher in responsive neurons than in non-responsive neurons (**Fig. 2F**; responsive: 7.5 Hz; non-responsive: 2.0 Hz; *p* = 4.5 × 10^-28^, Wilcoxon’s rank-sum test). The spike duration of responsive neurons was shorter than that of non-responsive neurons (**Fig. 2G**; responsive: 0.27 ms; non-responsive: 0.30 ms; *p* = 0.0012, Wilcoxon’s rank-sum test). These results suggest that each cluster, especially cluster #5, corresponds to a specific cell type in the SC.

Given the complexity of the responsiveness of the eight SC neuron clusters, a simpler analysis approach to examine the cluster properties was warranted. We initially noted that the response patterns could be broadly divided into excitatory-dominant (clusters #1, #2, #3, and #4) and inhibitory-dominant (clusters #5, #6, #7, and #8) patterns (**Fig. 2A**). Furthermore, we found that these multiphasic responses comprised three periods: **period 1** with fast transient excitation (clusters #1, #2, #3, and #5; blue-shaded area in **Fig. 2C**); **period 2** with slow inhibition (clusters #5, #6, and #7; purple-shaded area in **Fig. 2C**); and **period 3** with slow excitation (clusters #5, #6, and #7; orange-shaded area in **Fig. 2C**). The fast transient excitation in period 1 was often observed in clusters #1, #2, #3, and #5 neurons, which indicated that most SC responsive neurons receive direct monosynaptic input from the cortex. By contrast, the slow inhibition and excitation in period 2 were observed in clusters #4, #5, #6, #7, and #8 neurons and thus may be evoked by inputs from subcortical regions. No neurons showed a transient inhibition pattern, which is consistent with the fact that cortical projections are always excitatory.

### Comparison of single SC neuron responses by layer, ML, and AP locations

Previous anatomical studies have suggested that neuronal responses differ depending on location within the SC. We compared the proportion of responsive neurons by layer (**Fig. 3A**), ML zone (**Fig.3B**), and AP portion (**Fig. 3C**). The proportion of responsive neurons relative to all recorded neurons was higher in layers ventral to op than in Zo and SuG layers (**Fig. 3A**) and in the centrolateral and lateral SC than in the centromedial SC (**Fig. 3B**). There were no significant differences in the proportion of responsive neurons based on the AP axis (**Fig. 3C**). To investigate the recording location-dependency of the temporal activity patterns, we compared the occupancy of neurons in each cluster along the three axes. Neurons in clusters #1 and #2 were found more commonly in the op and InG layers than in the InWh and DpG layers (**Fig.3D**), in the centrolateral zone than in the lateral zone (**Fig. 3E**), and in the posterior portion than in the anterior portion (**Fig. 3F**). By contrast, neurons in clusters #6, #7, and #8 were observed more frequently in the InWh and DpG layers than in the op and InG layers (**Fig. 3D**), in the lateral zone than in the centrolateral zone (**Fig. 3E**), and in the anterior portion than in the posterior portion (**Fig. 3F**). Thus, the location-dependency of the response dynamics of SC neurons was evident; deeper, more lateral, more anterior SC neurons show longer and more multiphasic responses to cortical stimulation.

**Figure 3.**
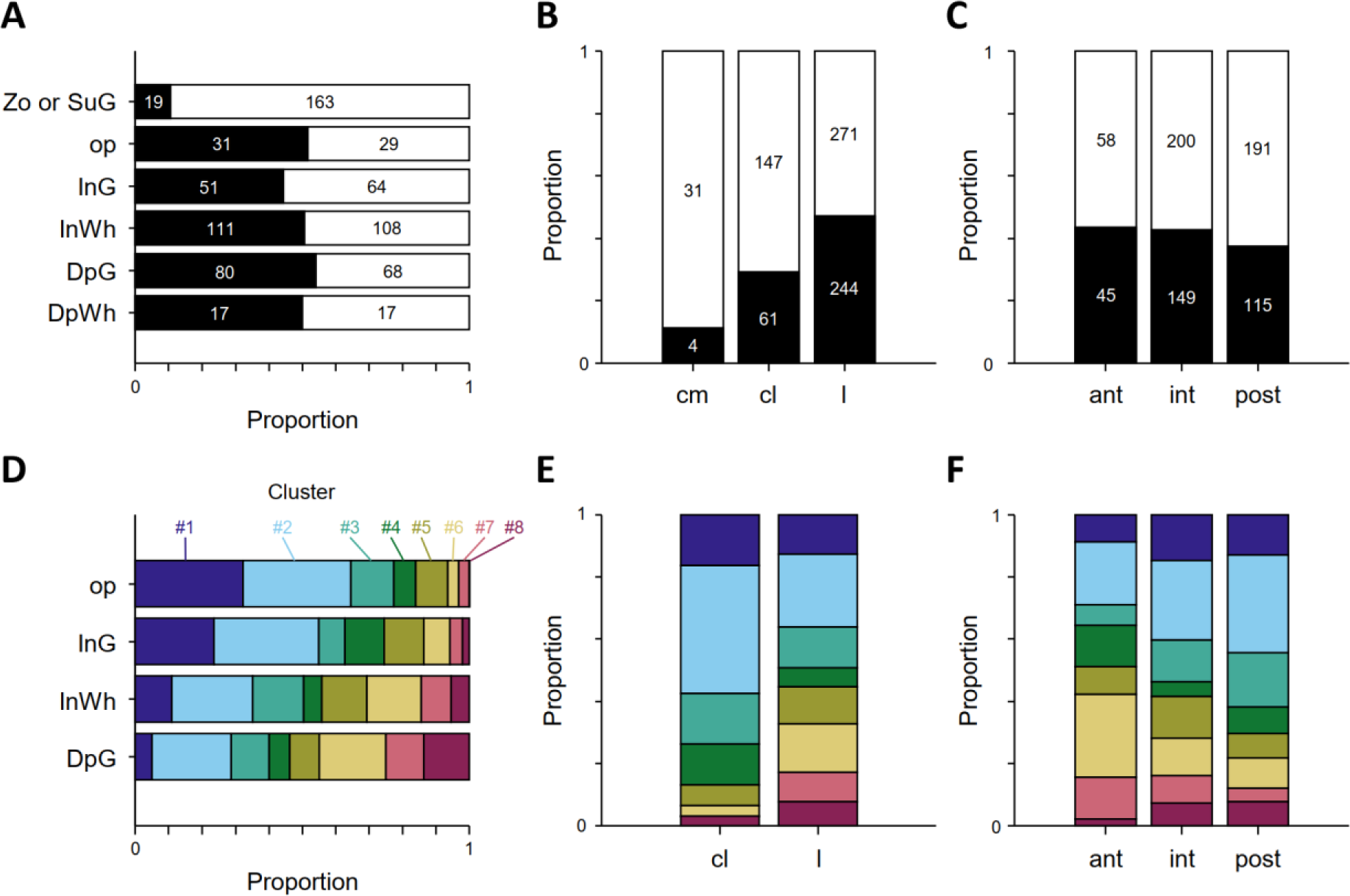
Distribution of recorded activity patterns. (A–C) The density and number of responsive neurons (black) and non-responsive neurons (white) recorded in: each layer (A), zones divided medially and laterally (B), and portions divided anteriorly and posteriorly (C). (D–F) The density of neurons classified into each cluster recorded in: each layer (D), zones divided medially and laterally (E), and portions divided anteriorly and posteriorly (F).

To examine the three-dimensional distribution of the degree of cortical-input integration, we defined the response duration, excitatory response duration, inhibitory response duration, and number of response phases (i.e., phase count) for each responsive neuron (**Methods**). There were significant differences between clusters in total duration (*p* = 0.007, ANOVA), excitatory duration (*p* = 8.4 × 10^-31^, ANOVA), inhibitory duration (*p* = 3.3 × 10^-11^, ANOVA), and phase count (*p* = 2.3 × 10^-15^, Kruskal-Wallis test; **Fig. 4A**). In clusters #5 and #6, neurons that showed excitatory responses lasting at least 50 ms accounted for 60.6% (20/33) and 80.5% (33/41), respectively. In cluster #8, neurons with inhibitory responses lasting at least 50 ms accounted for 81.0% (17/21), and in cluster #5, neurons with triphasic responses accounted for 75.8% (25/33). These findings demonstrate that these parameters captured the different dynamics of each cluster.

**Figure 4.**
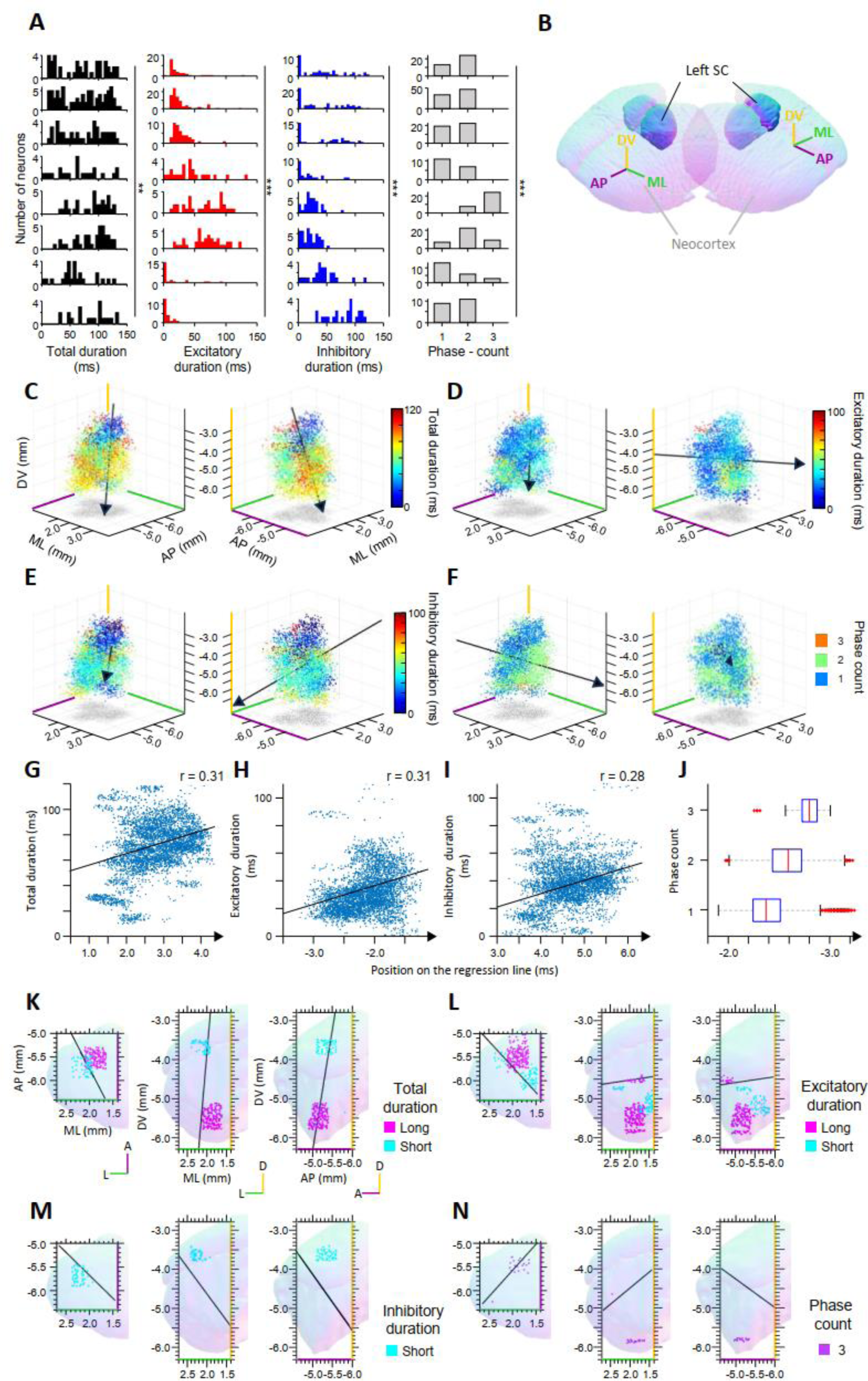
Three-dimensional distribution of the response complexity and duration of SC neurons. (A) Histograms of the excitatory duration (left), inhibitory duration (middle), and phase count (right) of neurons classified into each cluster. (B) The orientation of the neocortex and SC is displayed for the purpose of allowing for easier recognition of panel C–F. Three-dimensional representations were generated using an online software https://tfiers.github.io/3D-rat-brain/. Colors of the three axes correspond to those in (C–F). (C–F) Three-dimensional spatial plot of total duration (C), excitatory duration (D), inhibitory duration (E), and phase count (F) in the left SC. (G–J) Total duration (G), excitatory duration (H), inhibitory duration (I), and phase count (J) as a function of position on the regression line. The black lines correspond to the regression lines. Spearman’s correlation coefficient, r, was shown. (K-N) Location of neurons with significantly long or short total response durations (K), significantly long or short excitatory durations (L), significantly long or short inhibitory durations (M), and where triphasic response patterns were significantly more commonly recorded than the other voxels (N). SC, superior colliculus; DV, dorsoventral; ML, mediolateral; AP, anteroposterior

To visualize the three-dimensional distribution of these parameters, we plotted them in a 500 × 500 × 500 μm^3^ voxel using a color code (**Fig. 4C–F**). To show the gradient of these parameters within the SC, the regression line was overlapped in each plot. (black arrows, **Fig.4C**—F). Total duration was significantly positively correlated with the DV axis (**Fig.4C**; total duration: ML, *p* = 0.36; AP, *p* = 0.13; DV, *p* < 1.0×10^-10^; F-test). Inhibitory duration and excitatory duration were significantly positively correlated with the position along all three axes (**Fig.4D,E**; excitatory duration: ML, *p* < 1.0×10^-10^; AP, *p* < 1.0×10^-10^; DV, *p* = 9.6×10^-4^; inhibitory duration: ML, *p* < 1.0×10^-10^; AP, *p* = 2.2×10^-9^; DV, *p* < 1.0×10^-10^; F-test). Phase count was also positively correlated with all three axes (**Fig.4F**; ML, *p* < 1.0×10^-10^; AP, *p* < 1.0×10^-10^; DV, *p* < 1.0×10^-10^; F-test). Additionally, we analyzed the significance of the parameters in each voxel by comparing them with shuffled data (**Fig. 4K–N**, bootstrap test, see **Methods**). Voxels with significantly longer total durations were located 1.7–2.1 mm lateral, 5.4–5.8 mm posterior to the bregma, and 5.2–5.8 mm from the pial surface, corresponding to the intermediate portion of lateral zone in InWh or DpG layers. Voxels with significantly shorter total durations ere located 2.0–2.4 mm lateral, 5.6–6.0 mm posterior to the bregma, and 3.6–3.9 mm from the pial surface, corresponding to the intermediate portion of centrolateral or lateral zone in op, InG, or InWh layers (**Fig.4K**). Voxels with significantly longer excitatory durations were located 1.7–2.1 mm lateral, 5.0–5.9 mm posterior to the bregma, and 3.0–6.0 mm from the pial surface, corresponding to the anterior or intermediate portions of the lateral zone in the DpG layer (**Fig.4L**). Voxels with significantly shorter inhibitory durations were almost exclusively 2.0–2.4 mm lateral, 5.6–6.4 mm posterior to the bregma, and 3.5–3.9 mm from the pial surface corresponding to the intermediate or posterior portions of the centrolateral or lateral zones in the op, InG, and InWh layers (**Fig.4M**). Voxels with a higher occurrence of triphasic responses were found 1.7–2.1 mm lateral, 5.4–5.7 mm posterior, and 5.9 mm from the cortex surface, in the intermediate lateral zone in the DpG layer (**Fig. 4N**). Overall, neurons with long complex activity patterns were in the anterior portion of the lateral SC in the deep layers, whereas those with short simple activity patterns were concentrated in the posterior portion of the op and intermediate layers.

### Cortical area-dependent response patterns in the SC

Because we detected clear differences in response patterns within the SC depending on recording location, we next examined whether response patterns varied according to stimulus location in the cortex. To visualize the stimulus location-dependency of the response pattern, we mapped the excitatory duration, inhibitory duration, and phase count of each neuron onto the cortical surface where photostimulation induced the neuronal response (**Fig. 5A**). We observed clear cortical gradients on the map of both excitatory and inhibitory durations. Notably, the cortical area that caused long excitation and inhibition roughly corresponded to the rostral and caudal forelimb areas, respectively (Saiki et al. 2018; Hira, Ohkubo, Tanaka, et al. 2013; Hira, Ohkubo, Ozawa, et al. 2013) (**Fig. 5B**). This suggests that the SC calculates the difference between M1 and M2 activity to control forelimb movement.

**Figure 5.**
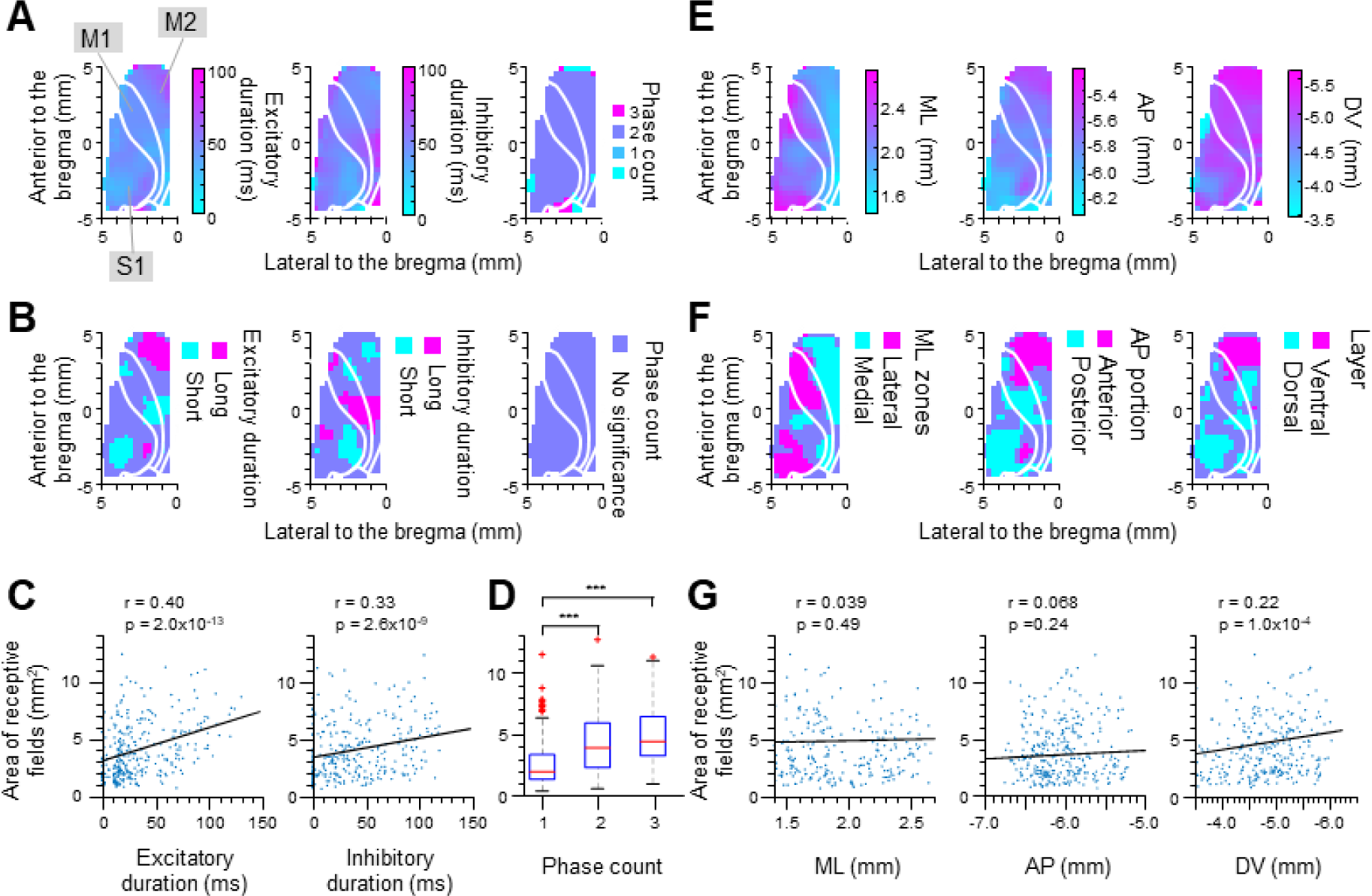
The relationship between the cortical areas and the locations and activity patterns of the SC neurons to which they project. (A) The excitatory duration (left), inhibitory duration (middle), and phase count (right) of the response induced via photostimulation was averaged for each 3 × 3 stimulus point on the cortex, and the corresponding center point on the cortex was color-coded accordingly. The color on the cortex indicates the temporal response pattern induced by the stimulus point in the responsive SC neurons. For visualization, the 3 × 3 surrounding points were averaged. (B) Similar analysis as in (A), but only significant points (*p* < 0.005, bootstrap test) are color-coded. (C) The area of the receptive field plotted as a function of excitatory (left) and inhibitory duration (right). The black lines correspond to the regression lines. Spearman’s correlation coefficient, r, and its *p*-value are shown. (D) Box plots of the receptive field area for neurons with monophasic, biphasic, or triphasic responses. \*\*\**p* < 0.001, Wilcoxon’s rank-sum test. (E) The same analysis as in (A), but plots show the mean location of the recorded neurons along the mediolateral (left), anteroposterior (middle), and dorsoventral axes (right). (F) Similar analysis as in (E), but only the significant points (*p* < 0.01, bootstrap test) are color-coded (G). The area of the receptive field plotted as a function of the mediolateral (left), anteroposterior (middle), and dorsoventral coordinates (right). Spearman’s correlation coefficient, r, and its *p*-value are shown. ML, mediolateral; AP, anteroposterior; DV, dorsoventral

We then compared the area of the receptive field and the excitatory duration, inhibitory duration, and phase count of the responsive neurons (**Fig. 5C,D**). We found that neurons with longer excitatory durations or those with longer inhibitory durations had a larger area of receptive fields. Additionally, the area of receptive fields was larger in neurons with biphasic or triphasic responses than in those with monophasic responses (*p* = 1.1×10^-8,^ ANOVA, monophasic vs biphasic, *p* = 4.26×10^-8^, monophasic vs triphasic, *p* = 3.36×10^-8^, Wilcoxon’s rank-sum test). These data indicate that neurons with long and multiphasic responses integrate signals from larger cortical areas than those with short and simple responses.

To visualize the relationship between cortical topography and SC locations, we mapped the locations of recorded neurons along the ML, AP, and DV axes on the cortical surface where photostimulation caused the neuronal response (**Fig. 5E**). We found cortical gradients all along the ML, AP, and DV axes. In all three maps, there were gradients from M2 to M1, where stimulation of M2 caused a neural response in the lateral, anterior, and ventral regions, whereas stimulation of M1 caused a neural response in the medial, posterior, and dorsal region (**Fig. 5F**). In addition, although the area of the receptive field did not significantly correlate with the location of recorded neurons along the ML or AP axis, neurons recorded in ventral regions had a significantly larger area of receptive fields (**Fig. 5G**). Taken together, these data indicate that SC neurons in the lateral, anterior, and ventral regions integrate broad cortical signals centered on M2 over long time windows and thus provide complex multiphasic outputs.

### Internal circuit in the intermediate portion of the lateral zone in the deep layer

Next, we examined what caused the difference in dynamic response patterns to cortical stimuli in SC neurons. The internal circuit within the SC is one candidate for generating such location-dependent dynamics. For example, the intermediate and deep layers of the SC comprise feedback or feedforward inhibitory circuits (Essig, Hunt, and Felsen 2021; Villalobos et al. 2018). We examined the cross-correlogram of the SC neuron pairs and found three candidates for inhibitory synaptic connections in the intermediate portion of the lateral zone in the DpG layer (**Fig. 6**). **Figures 6A and B** show that neuron #1 suppressed neurons #2 and #3. Neuron #1 exhibited a slow monophasic excitation pattern, and the activity of neurons #2 (orange) and #3 (green) reduced as neuron #1 was excited, which suggested that the inhibition of neuron #1 shapes the response patterns of neurons #2 and #3. By contrast, although neuron #4 has an inhibitory effect on neuron #5, as shown in the cross-correlogram (**Fig. 6C**), excitation of neurons #4 and #5 occurred simultaneously, immediately after photostimulation (**Fig. 6D**). These data suggest that not all response patterns can be explained by the SC internal circuit. The involvement of both internal circuits and input from other brain regions, such as SNr/GPi or CN, must be considered.

**Figure 6.**
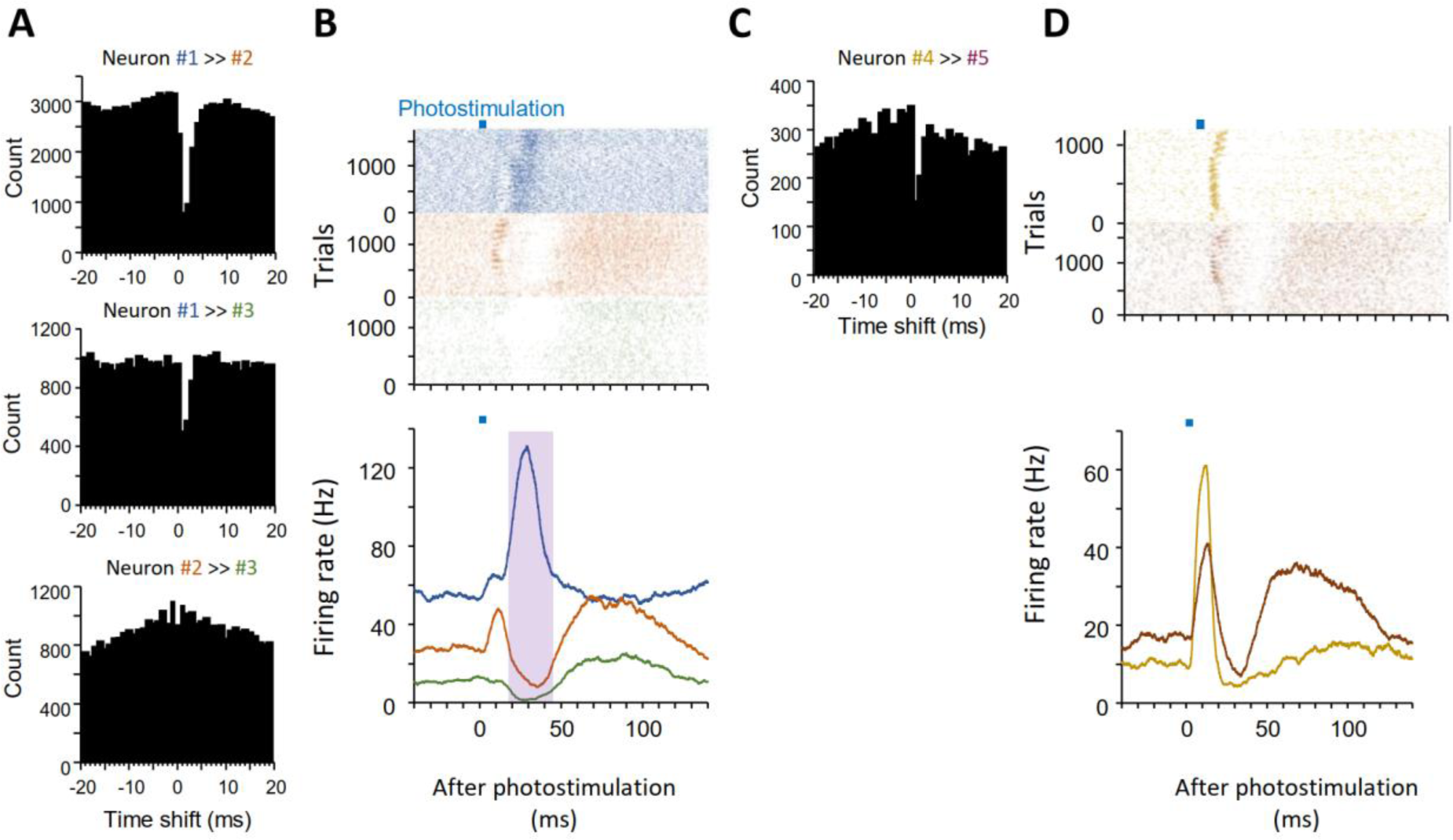
Cross-correlogram analysis and internal circuit in the SC. (A) Cross-correlogram between pairs of neurons #1, #2, and #3. (B) Raster plots and mean activity of neurons #1 (blue), #2 (orange), #3 (green). Purple-shaded area corresponds to period 2. (C) Cross-correlogram between neurons #4 and #5. (D) Raster plot and mean activity of neurons #4 (gold), and #5 (brown).

Anatomically confirmed projection sources that are considered candidates for inducing specific activity patterns in the SC are the basal ganglia output (i.e., the SNr) and cerebellar output (i.e., the CN). We re-examined the activity patterns of the SNr and CN in response to cortical stimuli using data acquired in a previous study (Yoshida et al. 2024) (**Fig. 7**). It was notable that the activity of both the SNr and CN was not ready for period 1 of the SC. However, the inhibitory inputs of period 2 could be influenced by the class 1 response of the CN and the class 2 response of the SNr. Specifically, the CN class 1 response showed an increase in activity after strong inhibition, which is consistent with the excitation after inhibition of SC clusters #5, #6, and #7. The period 2 excitatory inputs into SC neurons may be caused by CN class 2 responses and the inhibition of class 1 triphasic responses in the SNr. These results suggest that slow responses of the SC are affected by the CN and SNr. In particular, the inhibition of period 2 and the excitation of period 3, which are characteristic of the long and complex activity of the lateral SC, correspond to the CN class 1 response. The CN class 1 response is mediated by Purkinje cells, which suggests that the cerebellum has a considerable influence on the dynamic information processing of the lateral SC. Therefore, the cerebellum is a candidate region that heavily influences the dynamic information processing of the lateral SC.

**Figure 7.**
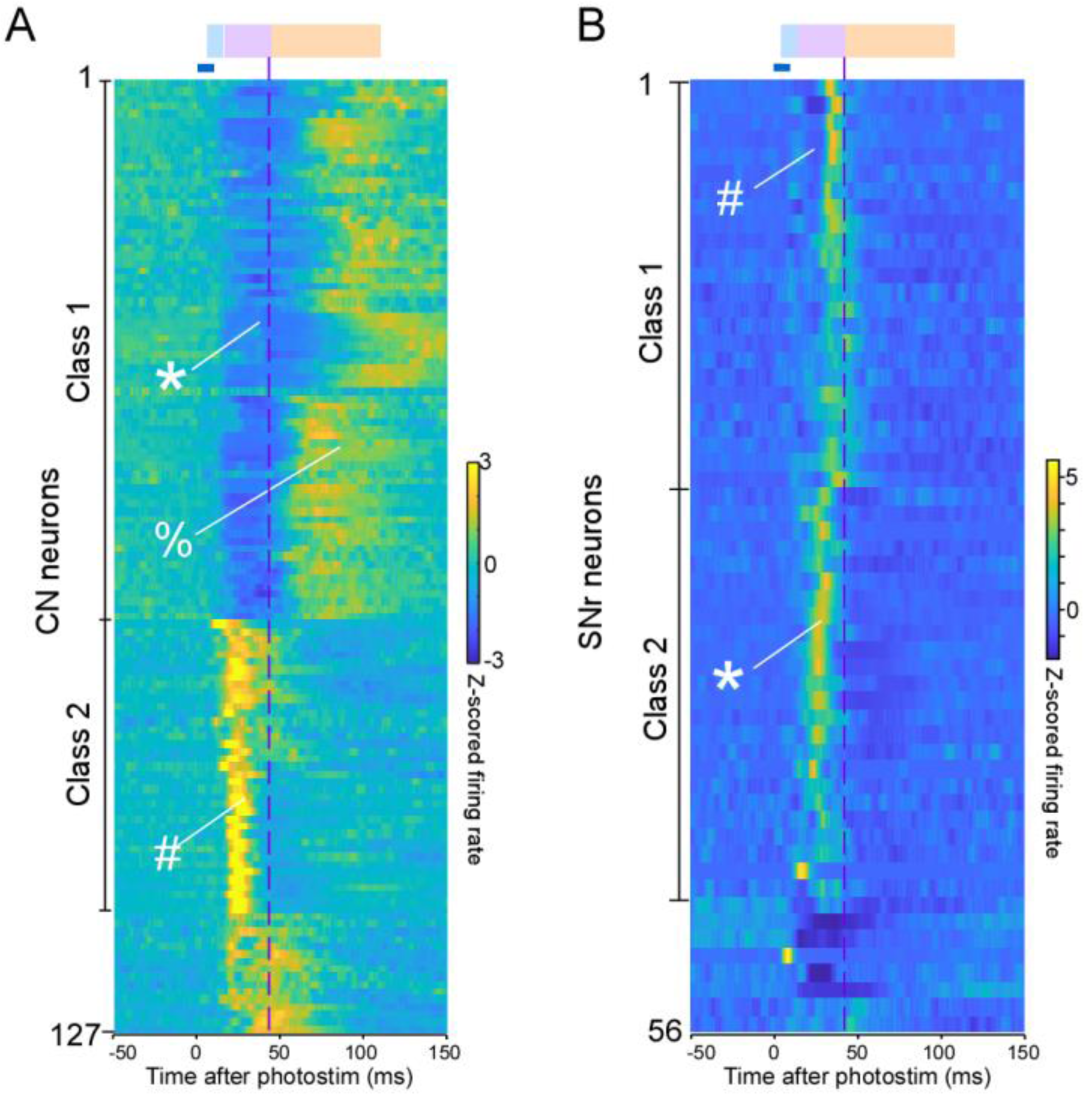
Responses of CN and SNr neurons to cortical stimulation. (A) Responsive neurons in the CN were sorted via K-means clustering, and class 1 and 2 responses were defined by the cluster. Inhibition (*) followed by excitation (%) is characteristic of a class 1 response, whereas a class 2 response is a simple transient excitation (#). (B) Responsive neurons in the SNr were sorted using K-means clustering, and class 1 and 2 responses were defined by the cluster. A triphasic response, which is composed of excitation-inhibition(#)-excitation, is characteristic of a class 1 response, whereas a class 2 response is a simple transient excitation (*). These figures were reproduced from figure 2d and figure 4d in our previous paper for explanation purppose (Yoshida et al. 2024).

### The SC deep layer projecting to the thalamic nuclei is connected to the cerebral cortex

The lateral and centrolateral SC activity may modulate higher functions by feeding information back to the cerebral cortex, especially the frontal cortex. The main thalamic nuclei that project to the frontal cortex are the VA-VL, VM, CL, and PF nuclei. To investigate which parts of the SC project to the thalamic nuclei, we injected AAVretro-CAG-tdTomato, which acts as a retrograde tracer, into these thalamic nuclei (**Fig. 8**). Upon injection into the VL nucleus, we noted that a few somas were visible in the SC. Following an injection into the VA, a few neurons were labeled in the ipsilateral SC. By contrast, when we injected the tracer into the VM or CL/PF, we found labeled neurons in layers InG and DpG of the lateral SC. These results are largely consistent with a previous study conducted in rodents (Benavidez et al. 2021; Yamasaki, Krauthamer, and Rhoades 1986; Krout et al. 2001; Horie et al. 2013). Together with the electrophysiological results, these data indicate that regions of the SC that exhibit complex and slow response patterns feed back to the frontal cortex via the ipsilateral thalamus, primarily the VM, CL, and PF. In addition, it is possible that projections to the intralaminar nuclei, such as the CL or PF, affect the cortical frontal cortex via the basal ganglia (Melleu and Canteras 2024).

**Figure 8.**
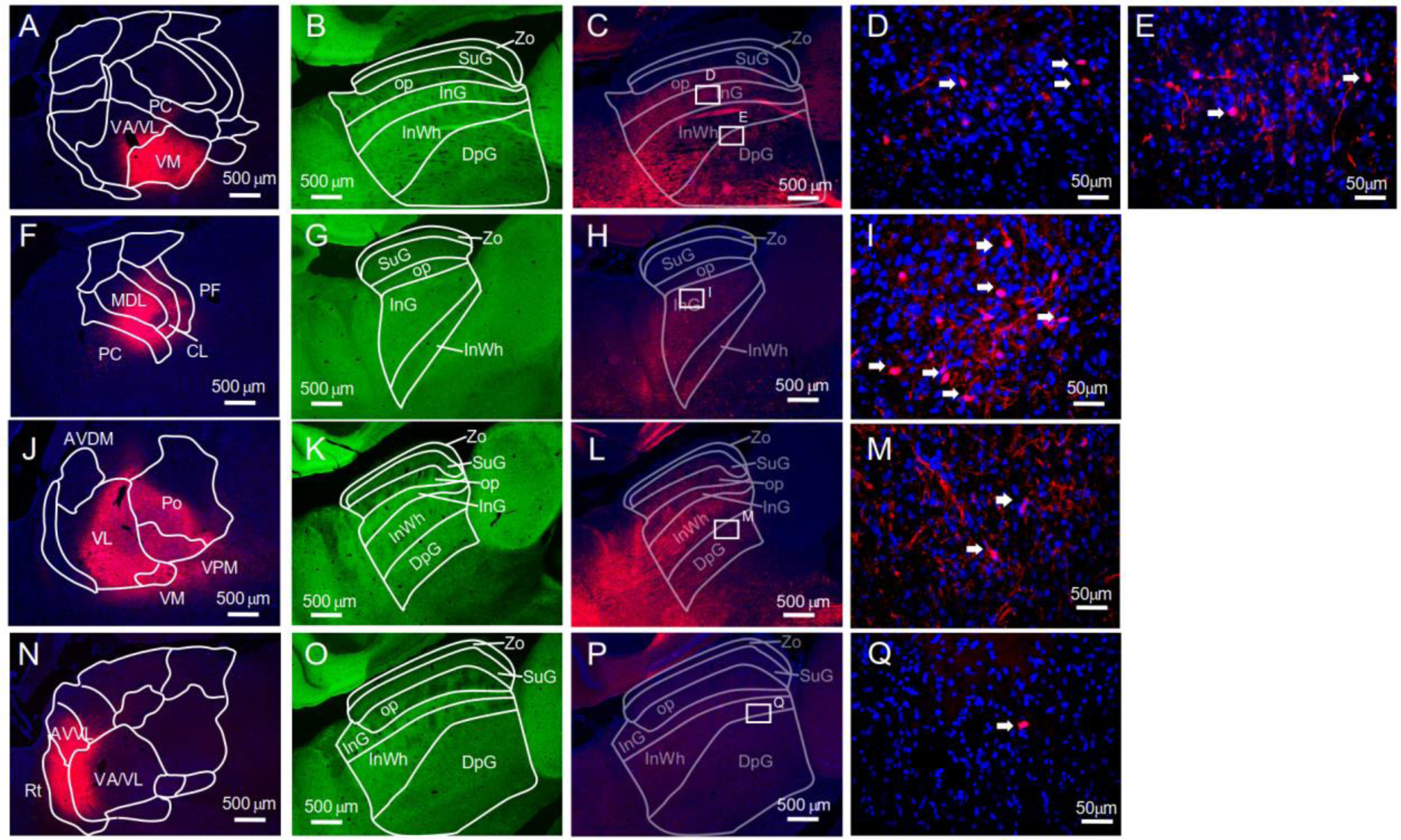
Retrograde tracing. **(A)** A sagittal fluorescence image of tdTomato (red) and DAPI (blue) at the injection site, VM. **(B)** A sagittal fluorescence image of Thy1-ChR2-Venus. Layer names are overlaid. **(C)** TdTomato (red) and DAPI (blue) images of the same locations shown in **(B**). **(D–E)** TdTomato (red) and DAPI (blue) images of the boxes specified in **(C). (F–I)** The same images as **(A–D)** for the CL injection. **(J–M)** The same images as **(A–D)** for the VL injection. **(N– Q)** The same images as **(A–D)** for the VA injection. Abbreviations: AVDM, anteroventral dorsomedial nucleus; AVVL, anterovental ventrolateral nucleus; CL, centrolateral nucleus; DpG, deep gray; InG, intermediate gray; InWh, intermediate white; MDL, mediodorsal nucleus lateral division; op, optic; PC, paracentral nucleus; PF, parafascicular nucleus; Po, posterior nucleus; Rt, reticular nucleus; SuG, superficial gray; VA, ventral anterior nucleus; VA/VL, ventral anterior/ventral lateral complex; VL, ventrolateral nucleus; VM, ventromedial nucleus; VPM, ventral posteromedial nucleus; Zo, zonal. The injection experiments for VM, VL, and VA were identical to those in our previous paper (Yoshida et al. 2024) with different projection areas analyzed.

## Discussion

In this study, we recorded single neuronal activity of the SC in response to cortical photostimulation. Neurons that showed slow inhibition, classified into clusters #6, #7, and #8, were more often recorded in the deep layer than in the intermediate layer, in the lateral zone than in the centrolateral zone, and in the anterior portion than in the posterior portion. By contrast, neurons that showed fast transient excitation, classified into clusters #1 and #2, were more often recorded in the op and intermediate layers than in the deep layer, in the centrolateral zone than in the lateral zone, and in the posterior portion than in the anterior portion. Therefore, there was a clear gradient in the dynamics along the depth, ML axis, and AP axis. These findings indicate that the SC has functionally distinctive structures that are defined by not only the input/output but also the dynamics induced by the cortex.

What specific inputs produce these different dynamics? Because internal synaptic circuits that affect response patterns are rarely found, differences between internal circuits within the SC may not be the key factor that contributes to different dynamics. Instead, based on the anatomical differences between the medial and lateral SC, differences in dynamics may be attributed to the information transmitted from the cerebellum and basal ganglia (J. Lee and Sabatini 2021; Rossi et al. 2016). Indeed, the major responses of the CN and SNr can affect period 2 of SC neuronal responses (**Fig. 7**). Thus, our results suggest that the diverse response dynamics of SC neurons are due to the signals sent from the other brain regions including cerebellum and basal ganglia.

How do the different dynamics in various SC regions impact other regions? The deep layer of the lateral SC projects to the PF, CL, and VM nuclei of the thalamus, which project to the frontal cortex (including M1 and M2), whereas the op and intermediate layers of the medial SC send axons to LP, which projects to higher-order visual areas (Blot et al. 2021; Murakami et al. 2022; Brenner et al. 2023). Therefore, the SC-LP pathway likely sends simple activity, similar to a relay-like operation, whereas the SC-PF/CL/VM pathway sends long and integrative signals via a dynamic circuit. This is in line with the idea that the medial SC processes ongoing and continuously changing visual information, whereas the lateral SC may integrates various information for action selection, decision-making, and top-down attention. It is also known that the lateral SC sends projections to the VTA and IO (May, Warren, and Kojima 2024; Solié et al. 2022), suggesting that the deep lateral SC dynamically controls learning in the basal ganglia and cerebellum. Aditionally, the deep lateral SC regulates forelimb movements via the medullary reticular nucleus and parvocellular reticular nucleus (Esposito, Capelli, and Arber 2014). By contrast, the medial and superficial SC induce escape behavior via the magnocellular reticular nucleus, cuneiform nucleus, or periaqueductal gray (Capelli et al. 2017; Caggiano et al. 2018; H. Wang et al. 2019; Evans et al. 2018; Campagner et al. 2023). Thus, the deep lateral SC is capable of regulating forelimb movements with complex dynamics directly via the brainstem or indirectly via the cerebral cortex, as well as modulating learning in the basal ganglia and cerebellum.

In our previous study (Yoshida et al. 2024), we found that the direct pathway of the basal ganglia and the mossy fiber-CN pathway synchronously can activate the thalamus 25–35 ms after cortical stimulation. Thus, we theorized that the cerebellum and basal ganglia would cooperatively drive the cerebral cortex via the thalamus approximately 30 ms after cortical activity. This time window was consistent with the inhibitory activity of clusters #5, #6, and #7 of SC neurons and the excitatory activity of cluster #4. Given that SC activity during this time window is largely inhibitory (inhibitory: 98/309, excitatory: 23/309), the SC (unlike the basal ganglia and cerebellum) likely coordinates decision-making and action selection by inhibiting the thalamus during this time window. Moreover, because considerable SC responses are activated earlier than this time window, the SC may affect the thalamus independently or competitively with the basal ganglia/cerebellar output. Additionally, the SC showed longer and more integrated dynamics in response to the photostimulation of M2 than to the photostimulation of M1 or S1; this was not observed in the CN or SNr. This supports the notion that the SC, in relation to the cortex, is independent of both the cerebellum and basal ganglia.

Recent analyses of SC cell types have elucidated cell type-specific functions (Brenner et al. 2023; Essig, Hunt, and Felsen 2021; Liu et al. 2022; Kaneda and Isa 2013; Doykos et al. 2020; Zingg et al. 2017; Li and Meister 2023; Farrow et al. 2019). Several of the eight clusters we identified may correspond to specific cell types. For example, cluster #4 neurons with short spike duration, which were excited exclusively during period 2, may represent a specific circuit element. The presence of seemingly unrelated functions, such as the output of ethologically relevant behaviors, decision-making and action selection, and regulation of basal ganglia and cerebellum learning signals, indicates that a specific cell type is responsible for a certain function and a certain dynamics. Our classification of responses to cortical stimuli will contribute to our understanding of the relationship between cell types and their functions.

## Acknowledgements

We thank T. Shimada, Reiko Hira for animal husbandry and genotyping, M. Kawabata, A. Rios, T. Sakairi, S. Aoki, M. Morishima for technical advice. This work was supported by JP22wm0525007 (RH), JP19dm0207089 (YI) from AMED, JP22H02731 (RH), JP20K22678 (RH), JP21B304 (RH), JP21H05134 (RH), JP21H05135 (RH), JP21H0524 2(YI), and JP23H02589 (YI) from MEXT/JSPS, JPMJCR1751 (YI) from JST, Nakatani Foundation (RH), Shimadzu Foundation (RH), Takeda Science Foundation (RH), The Precise Measurement Technology Promotion Foundation (RH), Tateishi Science and Technology Foundation (RH), and Research Foundation for OptoScience and Technology (RH).

## Author contributions

R.H, conceived the project. H.S. conducted electrophysiological experiments. S.T. conducted anatomical experiments. H.S. and T.Y. analyzed electrophysiological data. R.H. analyzed anatomical database. Y.I. supervised the project. H.S. and R.H. wrote the paper with the comments of all authors.

## Notes

### Competing Interest Statement

The authors have declared no competing interest.

